# HIT-EC: Trustworthy prediction of enzyme commission numbers using a hierarchical interpretable transformer

**DOI:** 10.1101/2025.02.01.635810

**Authors:** Louis Dumontet, So-Ra Han, Tae-Jin Oh, Mingon Kang

## Abstract

Accurate and trustworthy prediction of Enzyme Commission (EC) numbers is critical for understanding enzyme functions and their roles in biological processes. Despite the success of recently proposed deep learning-based models, there remain limitations, such as low performance in underrepresented EC numbers, lack of learning strategy with incomplete annotations, and limited interpretability. To address these challenges, we propose a novel hierarchical interpretable transformer model, HIT-EC, for trustworthy EC number prediction. HIT-EC employs a four-level transformer architecture that aligns with the hierarchical structure of EC numbers, and leverages both local and global dependencies within protein sequences for this multi-label classification task. We also propose a novel learning strategy to handle incomplete EC numbers. HIT-EC, as an evidential deep learning model, produces trustworthy predictions by providing domain-specific evidence through a biologically meaningful interpretation scheme. The predictive performance of HIT-EC was assessed by multiple experiments: a cross-validation with a large dataset, a validation with external data, and a species-based performance evaluation. HIT-EC showed statistically significant improvement in predictive performance when compared to the current state-of-the-art benchmark models. HIT-EC’s robust interpretability was further validated by identifying well-known conserved motifs and functional regions in the CYP106A2 enzyme family. HIT-EC would be a robust, interpretable, and reliable solution for EC number prediction, with significant implications for enzymology, drug discovery, and metabolic engineering. The open-source code is publicly available at: https://github.com/datax-lab/HIT-EC.

## 1 Introduction

The Enzyme Commission (EC) number classification system is a pivotal tool for categorizing enzymes based on the corresponding enzyme-catalyzed reactions [1–3]. An EC number consists of four hierarchical numbers (e.g., 1.1.1.1), which represent a class, a subclass, a sub-subclass, and a serial number, respectively [4]. For instance, the malate dehydrogenase enzymatic function is referred to as EC 1.1.1.37; the first level ‘1’ (EC ‘1’) denotes oxidoreductase; the second level (EC 1.’1’) indicates reaction in the CH-OH group of donors; the third number (EC 1.1.’1’) shows donor acceptors with NAD+ or NADP+; the last level (EC 1.1.1.’37’) refers to a malate dehydrogenase [5, 6]. This hierarchical classification characterizes enzyme functions and their roles in diverse biochemical pathways and facilitates the annotations of enzymes in biological databases. The significance of the EC number system extends beyond categorization, underpinning advancements in drug discovery, metabolic engineering, and ecological sustainability [7–9].

Accurate classification of EC numbers is essential for understanding enzyme functions and their roles in metabolic pathways. Computational approaches for EC number prediction are broadly categorized into three groups: (1) protein structure-based, (2) sequence similarity-based, and (3) machine learning-based methods. Protein structure-based models (e.g., COFACTOR [10], i-TASSER suite [11], AutoDock [12]) examine structural templates and perform homology analysis through protein threading using reference databases. While effective for identifying folds and functional sites, these models are constrained by the limited availability of high-quality structural data. Sequence similarity-based approaches (e.g., EFICAz [13], ModEnzA [14], PRIAM [15], EnzML [16]) utilize conserved patterns and motifs identified by multiple sequence alignment (MSA) to infer enzyme functions. Despite their substantial utility in identifying functional domains, these approaches (1) are computationally expensive, (2) are ineffective when confronted with the absence of closely related reference sequences, and (3) often require additional post-hoc analyses.

A promising solution is leveraging machine learning, which effectively captures complex relationships within enzyme sequences. The current state-of-the-art models include ECPred [17], DEEPre [18], DeepEC [19], ECPICK [20], DeepECtransformer [21], and CLEAN [22], which show enhanced predictive power when automating enzyme annotation. However, there still remains substantial room for improvement in their predictive performance, as these models often struggle with limited and imbalanced training data, particularly for enzymes with underrepresented functions where limited references are available [23]. Our study reveals that the state-of-the-art methods achieved an F1-score of only around 70% for underrepresented enzymes (e.g., *N* ≤ 25), which account for 41% of EC numbers in the dataset. This low performance highlights the challenges in achieving accurate predictions for underrepresented EC numbers.

Model interpretability also remains a critical challenge, particularly in establishing trustworthy predictions. Beyond achieving high predictive accuracy, it is essential to understand the rationale behind a model’s predictions to ensure biological relevance, especially when applied to high-stakes decisions. In EC number prediction, an interpretation scheme validates a model’s predictions by comparing highlighted amino acids with established biological knowledge, such as conserved motifs or functional sites, to ensure that the final prediction relies on biologically known and relevant components. In ECPICK, model interpretation provided insights into how specific motifs or regions influence predictions, yielding domain-specific evidence [20]. In addition to increasing trustworthiness in the predictions, it also has the potential to uncover new motif sites or functional regions, which could drive further experimental investigations and expand our understanding of enzyme functions.

Incomplete annotations in biological databases present a significant opportunity to improve the predictive performance of deep learning models. For instance, as of September 2022, 15.4% of protein sequences in the Swiss-Prot database are incompletely annotated, often missing last levels of the EC classification (e.g., 1.1.-.-). These incomplete annotations arise from several factors, including the labor-intensive and time-consuming nature of experimental characterization, the vast number of newly discovered enzymes that have not yet been studied, and the challenges in reliably assigning EC numbers to enzymes with novel or ambiguous functions [23]. These limitations address the potential to enhance model performance by developing methods that can effectively handle incomplete annotations. To the best of our knowledge, all of the current state-of-the-art models rely on only completely annotated data when training for EC number prediction, which deprives these models of valuable data.

In this study, we develop a novel Hierarchical Interpretable Transformer model (HIT-EC) that advances EC number classification by leveraging the hierarchical structure of EC numbers and incompletely labeled sequences (Fig. 1). HIT-EC employs a four-level transformer architecture that aligns with the EC number hierarchy, which provides context-aware predictions at each level. The major contributions of the HIT-EC model are as follows: (1) statistically significant improvement of the predictive performance, (2) a novel training strategy to handle incomplete annotations, and an evidential approach to provide trustworthy predictions (Fig. 1). The performance of HIT-EC was assessed by extensive experiments across multiple evaluation settings with a large dataset (over 200,000 sequences), including (1) cross-validation experiments, (2) external validation using newly registered enzymes, and (3) the evaluation of the predictive performance on complete genomes of various species. In the experiments, HIT-EC consistently outperformed the state-of-the-art methods across all evaluation settings. HIT-EC improved micro- and macro-averaged F1-scores by at least 6% and 4%, respectively, in the cross-validation experiment, and also showed a 6% improvement in F1-score for underrepresented enzyme classes with fewer than 25 sequences. Furthermore, we demonstrated the trustworthiness of HIT-EC’s predictions by comparing the regions highlighted by our method to established biological knowledge within the CYP106A2 enzyme family, which provided evidence of the similarities between our interpretation and well-known functional and structural characteristics.

**Fig. 1.**
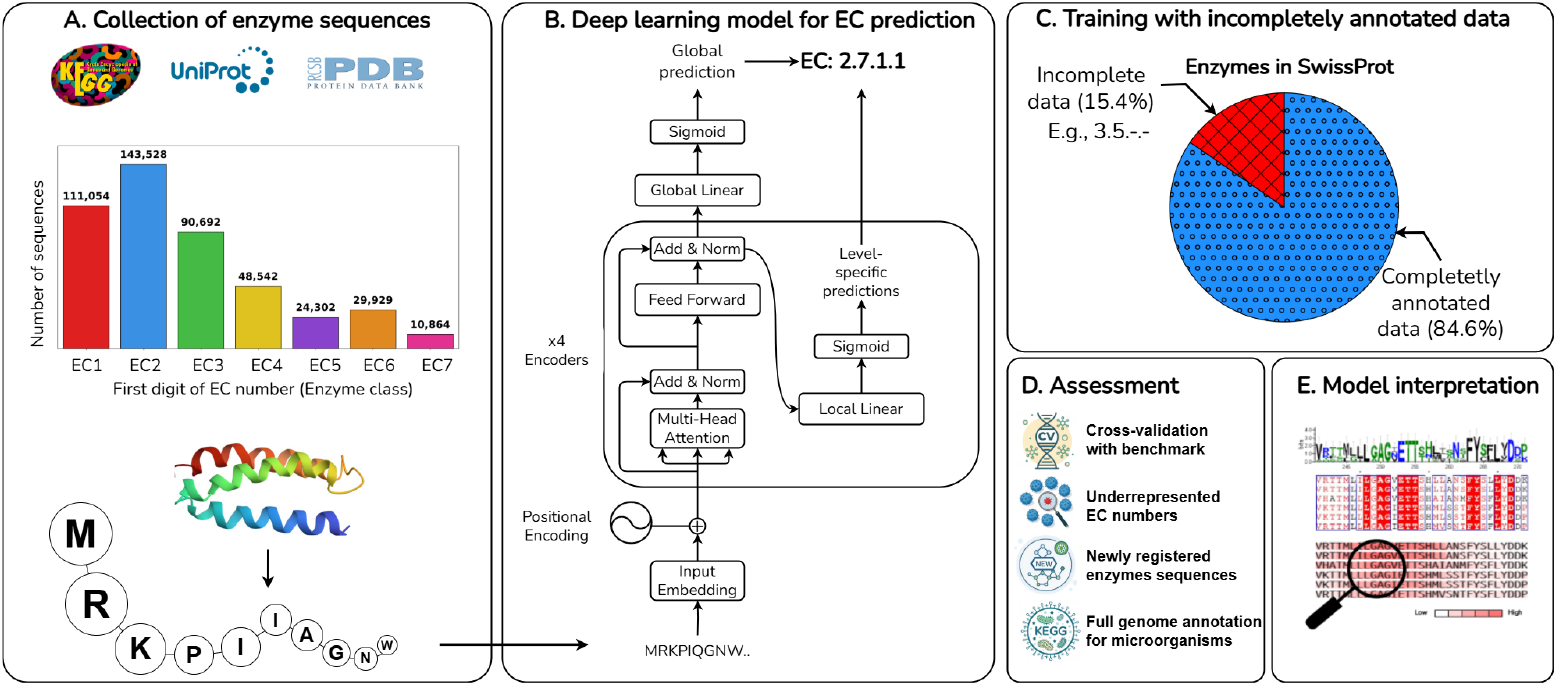
An overview of the study. (A) Collection of protein sequences from Uniprot, the Protein Data Bank, and the KEGG database. (B) Innovative hierarchical interpretable using both local and global flows named HIT-EC. (C) HIT-EC trains with incomplete EC numbers with the masked loss mechanism. (D) Assessment of the predictive performance against three state-of-the-art benchmark models across various experiment settings. (E) HIT-EC provides trustworthy predictions validated by aligning the model’s interpretation with established biological knowledge.

## 2 Methods

In this section, we elucidate the proposed HIT-EC model, focusing on three core components: the architecture of the proposed framework, the approach for training with incomplete EC annotations, and the strategy for enhancing model interpretability.

### 2.1 Architecture of HIT-EC

HIT-EC consists of five modules: (1) an embedding layer that encodes a protein sequence to characterize its physicochemical properties as numerical representations, (2) a positional encoding layer that incorporates sequence order information, (3) four transformer encoders, each of which corresponds to a level of the EC hierarchy and estimates level-specific *local* predictions, (4) a global linear layer that generates a *global* prediction of the EC numbers, and (5) the aggregation of the global prediction and the level-specific local predictions to form the final prediction. The HIT-EC architecture captures both local and global dependencies within sequences, while leveraging the hierarchical structure of the EC nomenclature.

First, the embedding layer converts amino acid sequences into physicochemical representations [24]. Each amino acid in a sequence is mapped to a fixed-dimensional vector using an embedding matrix **E** ∈ ℜ^23*×d*^, where *d* is the dimension of the embeddings. The value 23 corresponds to the 20 amino acids, the special codon ‘X’ (representing the amino acids ‘B’, ‘Z’, ‘U’, and ‘O’), the classification token, and the padding token. Each row of **E** represents a specific amino acid, and the matrix values are learned in the training phase to capture meaningful biochemical and structural properties. The embedding layer projects each sequence into a continuous vector space. By transforming the discrete sequence data into a numerical format, the embedding layer provides the foundation for downstream layers to process and extract relevant features for enzyme classification.

Second, the positional encoding layer integrates sequence order information into the model to capture the sequential nature of protein structures [25]. Since transformers lack inherent sequence-order awareness, positional encodings are added to the input embeddings to provide this positional context. HIT-EC employs sinusoidal positional encodings, where each position (i.e., *p*) in the sequence is assigned a unique vector using sine and cosine functions of different frequencies. The positional encoding (i.e., P(·)) for each dimension *i* at the position *p* is computed as:

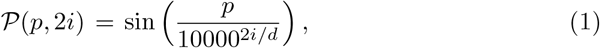

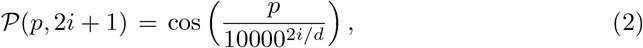

where 2*i* and 2*i* + 1 represent even and odd indices, respectively. These encodings are then added to the amino acid embeddings, resulting in **𝒜** ∈ ℜ^*S×d*^, where *S* is the sequence length (i.e., *S* = 1, 024). Each row of **𝒜** corresponds to an amino acid, containing both physicochemical properties and positional information. This positional information is crucial for capturing sequence-specific patterns, such as conserved motifs and functional regions, which are associated with the EC number classification.

Third, HIT-EC introduces a novel transformer-based architecture to implement the complementary paradigms of the local and global flows to enhance the hierarchical classification of enzyme functions. The local flow captures the dependencies between each level and the previous level of the EC hierarchy to capture hierarchical relationships [26]. This ensures that the predictions for each level are informed by the previous level’s output. On the other hand, the global flow treats each level independently by reintroducing the original sequence embeddings into the subsequent encoders:

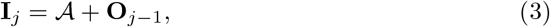

where **I**_*j*_ is the input of the *j*^*th*^ encoder, and **O**_*j*−1_ is the output of the *j* − 1^*th*^ encoder. Note that **I**_1_ = **𝒜**. To enable global classification for enzyme prediction, we add a special classification token as the first token of the input sequence. This token serves as a summary representation of the entire sequence. By including this token, the model can aggregate sequence-level information efficiently for downstream classification tasks. By combining both local and global flows, HIT-EC considers both the specific relationships at each hierarchical level and the overall context of the protein sequence.

HIT-EC includes a hierarchical structure of four encoders to predict each level of the EC hierarchy, where each encoder is composed of a multi-head self-attention mechanism and a position-wise feed-forward neural network. The multi-head self-attention mechanism captures relationships between amino acids within a sequence, identifying multiple relevant interactions in parallel. The position-wise feed-forward network refines the representations learned by the attention mechanism to detect more complex patterns. Layer normalization and residual connections are also incorporated to stabilize training and enhance convergence. Each encoder generates both a sequence representation and a level-specific prediction.

Fourth, HIT-EC simultaneously generates global predictions for all levels of the EC hierarchy using a linear layer after processing through the encoders. The input of this layer is the representation of the classification token from the fourth encoder.

Finally, the final prediction is produced by aggregating the local (*P*_ℒ_) and global (*P*_𝒢_) predictions:

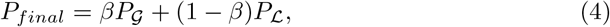

where *β* ∈ [0, 1] is a hyperparameter that determines the balance between the hierarchical specificity provided by the local predictions with the comprehensive view captured by the global prediction.

### 2.2 Training with incomplete EC number data

HIT-EC incorporates a novel masked objective function to handle incompletely labeled data in hierarchical multi-label classification settings. Specifically, the masked loss function adjusts the contribution of unannotated levels in the EC hierarchy, such that only the annotated levels contribute to the computed loss during training. This approach utilizes binary cross-entropy (BCE) loss to quantify the discrepancy between predicted and ground truth values. Let *y*_*ij*_ be the label for the sequence *i* at level *j*. We define a binary mask **M**, with *m*_*ij*_ = 1 if the ground truth *y*_*ij*_ is available and *m*_*ij*_ = 0 otherwise. The total loss for a batch of N sequences across the four levels of the EC hierarchy is computed as:

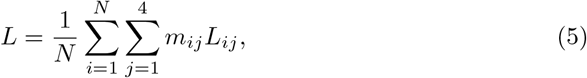

where *L*_*ij*_ is the BCE loss for the predicted probability *ŷ*_*ij*_ at level *j* of the EC hierarchy, given the ground truth *y*_*ij*_.

### 2.3 Model interpretation for domain-specific evidence

HIT-EC’s interpretation scheme integrates attention flow with gradient-based relevance propagation, offering a comprehensive view of the model’s decision-making process. HIT-EC identifies biologically significant regions in protein sequences, such as conserved motifs or functional domains like active sites or binding pockets, that contribute to enzymatic activity. Additionally, it accommodates multi-label classification, allowing for separate relevance scores to be assigned for each level of the EC number prediction, thereby providing independent explanations for each hierarchical level.

Relevance scores are computed for each amino acid by adapting an explainable framework for transformer-based models [27]. The method propagates gradients of the output prediction with respect to the input tokens through the attention layers and combines them with the attention weights to generate the final relevance scores. The contribution of each input feature is appropriately reflected in the corresponding relevance score. The relevance score *r*_*i*_ for an input token *x*_*i*_ is computed as:

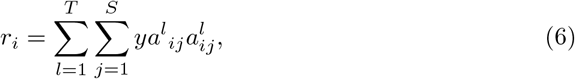

where *T* is the number of attention layers (i.e., *T* = 4), *S* is the number of tokens (i.e., *S* = 1, 024), *a*^*l*^_*ij*_ is the attention weight from token *i* to token *j* at layer *l*, and *y* is the model’s final prediction.

## 3 Results

In this section, we present the performance evaluation of HIT-EC through intensive experiments in various settings: (1) the micro- and macro-averaged F1 scores on the cross-validation dataset, (2) the performance of HIT-EC on underrepresented classes within this dataset, (3) the predictive performance on newly registered enzymes, and the performance on various species from the KEGG database [28].

### 3.1 Cross-validation for performance comparison

We evaluated the performance of HIT-EC by comparing it to state-of-the-art models in a cross-validation setting. We considered manually curated protein sequences from Swiss-Prot [29] and the Protein Data Bank (PDB) [30], released prior to September 2022. Sequences were filtered to retain only non-redundant entries with a maximum length of 1,023 amino acids. We excluded EC numbers associated with less than ten sequences. This data preprocessing resulted in the cross-validation dataset that includes approximately 200,000 sequences categorized into 1,938 EC numbers. The dataset was split into training (80%), validation (10%), and test (10%) sets by stratified sampling, preserving the class ratios, for the assessment. We considered three state-of-the-art benchmark methods, CLEAN [22], ECPICK [20], and DeepECtransformer (DeepECT) [21], for the comparison. For the hyper-parameter tuning, we used a framework for hyper-parameter optimization, Optuna [31], where the hyper-parameters were optimized to minimize loss on the validation set. For the final predictions, we selected class-specific thresholds which maximize the F1-score on the validation set for each EC number. The optimal hyper-parameter values of HIT-EC were an attention head count of 2, an embedding dimension (*d*) of 1,024, a dropout rate of 0.1, a *β* of 0.59, and a learning rate of 8.75e-5. For the benchmark models, we used the optimal hyper-parameters reported in their original papers. For training HIT-EC, the Stochastic Weight Averaging method was applied during the last 15 epochs to improve the model’s generalization performance and stability [32]. All models were trained on the same dataset, ensuring a consistent and fair comparison across all evaluation tasks. To ensure reproducibility, this experiment was conducted ten times.

HIT-EC achieved the highest micro-averaged F1-score of 0.93 ± 0.01, followed by CLEAN (0.88 ± 0.01), ECPICK (0.81 ± 0.07), and DeepECT (0.79 ± 0.05) (Fig. 2A). HIT-EC demonstrated an 18% improvement over DeepECT, a 15% improvement over ECPICK, and a 6% improvement over CLEAN. The statistical outperformance was assessed by Wilcoxon signed-rank tests (*p <* 0.01 for all comparisons). Note that CLEAN always predicts at least one EC number for any protein sequences, which showed relatively high F1-scores but can lead to a large number of false positives and a False Discovery Rate (FDR) of 1 with non-enzymes. In contrast, other models employ thresholding for the final predictions, which assign EC numbers for high confidence scores only. We considered any prediction failures due to low confidence as negatives in the evaluation. The FDRs computed using non-enzyme protein sequences with taxonomy information in Swiss-Prot were 0.15, 0.17, and 0.16 for DeepECT, ECPICK, and HIT-EC, respectively. However, the FDR of CLEAN was 1.

**Fig. 2.**
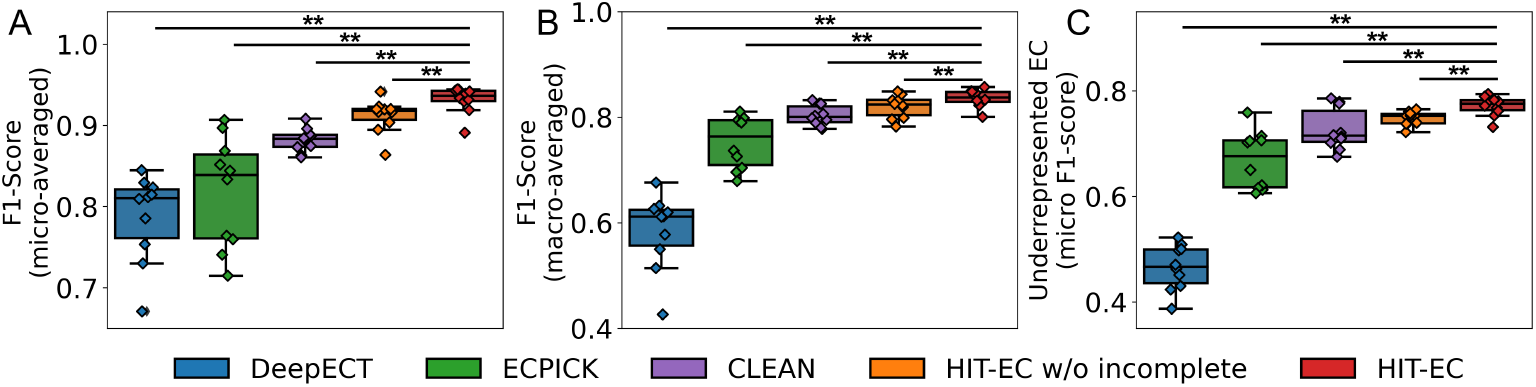
Performance comparison on the cross-validation dataset containing 200,000 sequences from Swiss-Prot and PDB. (A) Micro-averaged F1-scores over ten experiments, (B) macro-averaged F1-scores, and (C) micro-averaged F1-scores with only underrepresented EC classes (N≤25). HIT-EC showed statistically significant improvement by training incomplete data (p*<*0.01). ** indicates statistically significant improvement with p*<*0.01 using the Wilcoxon ranked signed test.

HIT-EC also showed the highest macro-averaged F1-score of 0.84 ± 0.02, surpassing CLEAN (0.80 ± 0.02), ECPICK (0.75 ± 0.05), and DeepECT (0.58 ± 0.07) (*p <* 0.01 for all comparisons) (Fig. 2B). The higher macro-averaged F1-score indicates HIT-EC’s robust performance across all EC numbers, while micro-averaged F1-scores can be biased towards the majority classes.

We further evaluated the performance considering only underrepresented EC classes, which contain fewer than 25 sequences. The cross-validation dataset included 799 underrepresented EC numbers. HIT-EC also demonstrated superior performance in this setting, achieving the highest micro-averaged F1-score of 0.77 ± 0.02, followed by CLEAN (0.73 ± 0.04), ECPICK (0.67 ± 0.05), and DeepECT (0.47 ± 0.04) (Fig. 2C). HIT-EC outperformed DeepECT by 64%, ECPICK by 15%, and CLEAN by 5% (*p <* 0.01 for all comparisons).

Furthermore, we examined HIT-EC’s performance with and without incompletely annotated data when training to assess the proposed learning strategy. The HIT-EC model that was trained with incomplete data showed an improvement of at least 2% over the model trained without incomplete data in the micro-/macro-averaged F1-scores and underrepresented EC classes (*p <* 0.01) (Fig. 2A-C).

### 3.2 External validation using newly registered enzymes

For the external validation, we conducted experiments using newly registered enzyme sequences in Swiss-Prot, referred to as the *New-392* dataset [22, 33]. This dataset consists of 392 protein sequences, released after September 2022, which ensures that none of these sequences were part of the training process for any of the models.

First, we considered the models trained in the cross-validation experiments, where ten optimal models were available for each benchmark method. We applied the models to the New-392 dataset and computed micro-averaged F1-scores. The models which were trained on the cross-validation dataset covered only 1,938 EC numbers. Thus, any sequences falling outside of this coverage were considered as negatives when computing the confusion matrix. Out of the 392 sequences, 192 were outside of the coverage in this experiment. HIT-EC produced the highest micro-averaged F1-score of 0.69±0.04, surpassing ECPICK by 3% (0.67 ± 0.09), DeepECT by 15% (0.60±0.05), and CLEAN by 44% (0.48 ± 0.07) (*p <* 0.01 for all comparisons) (Fig. 3A).

**Fig. 3.**
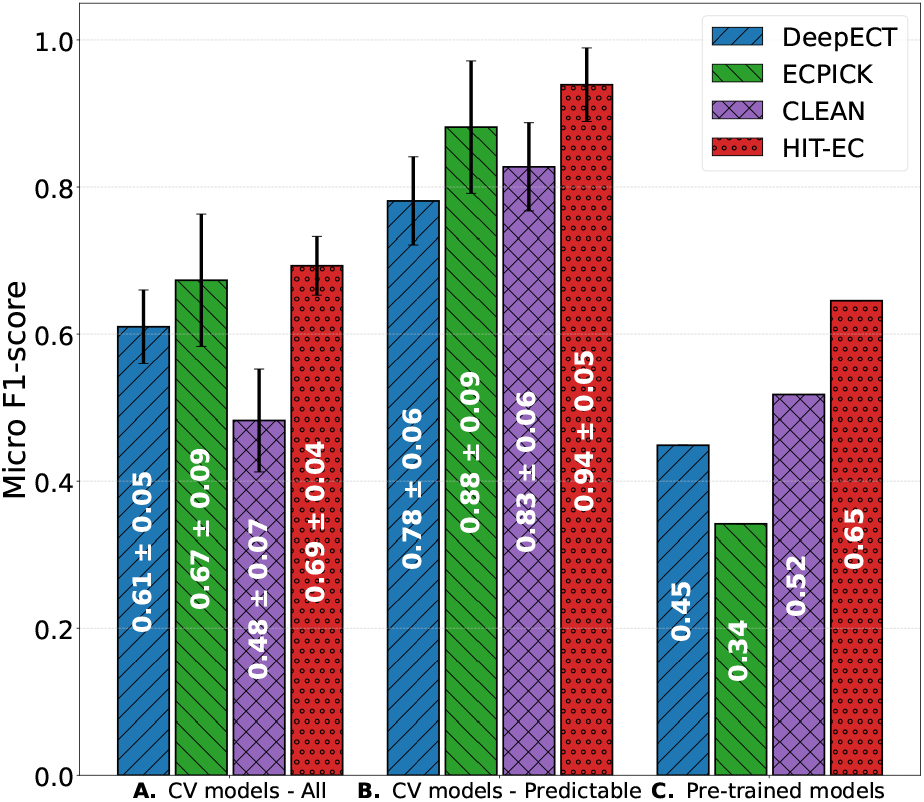
Performance comparison on the New-392 dataset for the external validation. (A) Cross-validation models using all protein sequences (N=392), (B) cross-validation models using only sequences with predictable EC numbers (N=200). (C) Performance using the pre-trained models that are publicly available.

In the second evaluation setting, we only considered sequences within the coverage of the cross-validation models. HIT-EC achieved a micro-averaged F1-score of 0.94 ± 0.05, significantly outperforming CLEAN by 7% (0.88 ± 0.06), ECPICK by 13% (0.83 ± 0.09), and DeepECT by 21% (0.78 ± 0.06). (*p <* 0.01 for all comparisons) (Fig. 3B). There were 15, 17, and 12 failures in the EC number assignments in ECPICK, DeepECT, and HIT-EC, respectively.

Additionally, we performed the experiments using the pre-trained models that are publicly available. We downloaded the latest version of the pre-trained models from GitHub. For HIT-EC, we re-trained the model using all available datasets (before September 2022), including Swiss-Prot, PDB, and manually curated entries from the KEGG database. To increase the coverage of HIT-EC’s predictable EC numbers, we incorporated enzymes from the TrEMBL database. For the EC numbers represented by fewer than 80 sequences, additional sequences were added from TrEMBL, where the EC numbers of the TrEMBL data were re-labeled by aligning these sequences with sequences in Swiss-Prot and PDB using DIAMOND (minimum percent identity of 50% and coverage of 75%). The augmentation produced around 450,000 sequences for the training dataset. The coverage of the resulting HIT-EC model was 4,255 EC numbers. HIT-EC still showed the highest micro-averaged F1-score of 0.65, followed by CLEAN (0.52), DeepECT (0.45), and ECPICK (0.34) (Fig. 3C). Note that the pre-trained models were trained with different datasets and various EC number coverages.

### 3.3 Species-specific performance comparison using KEGG

We assessed the models’ performances on complete genome datasets derived from KEGG to further evaluate the robustness of these models, using the benchmark models trained in the cross-validation experiments. In this experiment, we considered microbial genomes that exhibit a great diversity in sequence composition and evolutionary trajectories. Unlike human proteins which have been extensively studied and annotated, microbial proteomes often include novel or poorly characterized sequences. Specifically, we evaluated the models using fourteen microbial species, including Shigella flexneri (strain 301), Acinetobacter pittii (strain PHEA-2), Campylobacter jejuni (strain NCTC 11168), Staphylococcus aureus (strain NCTC 8325),

Pseudomonas aeruginosa (strain PAO1), Bacillus subtilis (strain 168), Escherichia coli (strains O157:H7 and K-12), Mycobacterium tuberculosis (strain H37Rv), Klebsiella pneumoniae (strain HS11286), Caulobacter vibrioides (strain NA1000), Chlamydia trachomatis (strain D/UW-3/CX), Listeria monocytogenes (strain EGD-e), and Coxiella burnetii (strain RSA 493).

HIT-EC exhibited the highest F1-scores in eight of the fourteen species and showed the best performance on average. HIT-EC achieved the micro-averaged F1-score of 0.81 ± 0.06 on average, which was significantly higher than ECPICK (0.78 ± 0.04), CLEAN (0.77 ± 0.04), and DeepECT (0.67 ± 0.08) (*p <* 0.05 for all comparisons) (Table 1). The overall superior performance of HIT-EC on this comprehensive genome dataset indicates its robustness and capability in large-scale enzyme function prediction. The experimental result also implies that HIT-EC effectively captures patterns in diverse protein families and is robust to variations in sequence composition.

**Table 1.** Performance Comparison across species using KEGG.

| Species (strains) | EC PICK | DeepECT | CLEAN | HIT-EC | Samples # |
| --- | --- | --- | --- | --- | --- |
| *Shigella flexneri* (strain 301) | 0.85 ± 0.12 | 0.79 ± 0.04 | 0.83 ± 0.01 | **<u>0.90 ± 4e-3\*\*</u>** | 1,346 |
| *Acinetobacter pittii* (strain PHEA-2) | **<u>0.76 ± 0.06\*\*</u>** | 0.63 ± 0.08 | 0.74 ± 0.02 | 0.73 ± 0.01 | 1,365 |
| *Campylobacter jejuni* (strain NCTC 11168) | 0.77 ± 0.04 | 0.57 ± 0.03 | **<u>0.81 ± 0.01\*\*</u>** | 0.79 ± 0.02 | 658 |
| *Staphylococcus aureus* (strain NCTC 8325) | 0.79 ± 0.06 | 0.64 ± 0.03 | 0.79 ± 0.01 | **<u>0.80 ± 0.01\*</u>** | 991 |
| *Pseudomonas aeruginosa* (strain PAO1) | 0.79 ± 0.07 | 0.62 ± 0.02 | 0.77 ± 0.02 | **<u>0.79 ± 0.01\*\*</u>** | 1,829 |
| *Bacillus subtilis* (strain 168) | 0.79 ± 0.12 | 0.78 ± 0.04 | 0.75 ± 3e-3 | **<u>0.87 ± 0.02\*\*</u>** | 1,370 |
| *Escherichia coli* (strain O157:H7) | 0.83 ± 0.03 | 0.78 ± 0.03 | 0.82 ± 0.01 | **<u>0.91 ± 0.01\*\*</u>** | 1,547 |
| *Escherichia coli* (strain K-12) | 0.83 ± 0.03 | 0.79 ± 0.03 | 0.82 ± 0.01 | **<u>0.91 ± 0.01\*\*</u>** | 1,579 |
| *Mycobacterium tuberculosis* (strain H37Rv) | 0.76 ± 0.06 | 0.61 ± 0.05 | 0.67 ± 0.03 | **<u>0.76 ± 0.01</u>** | 1,412 |
| *Klebsiella pneumoniae* (strain HS11286) | **<u>0.83 ± 0.14</u>** | 0.74 ± 0.01 | 0.77 ± 0.01 | 0.83 ± 0.01 | 1,907 |
| *Caulobacter vibrioides* (strain NA1000) | 0.70 ± 0.05 | 0.60 ± 0.09 | **<u>0.73 ± 0.01\*</u>** | 0.73 ± 0.01 | 1,907 |
| *Chlamydia trachomatis* (strain D/UW-3/CX) | 0.77 ± 0.04 | 0.55 ± 0.04 | **<u>0.78 ± 0.01\*\*</u>** | 0.76 ± 0.01 | 303 |
| *Listeria monocytogenes* (strain EGD-e) | 0.72 ± 0.10 | 0.72 ± 0.01 | 0.79 ± 0.03 | **<u>0.80 ± 0.01\*\*</u>** | 1,026 |
| *Coxiella burnetii* (strain RSA 493) | **<u>0.79 ± 0.03\*\*</u>** | 0.60 ± 0.03 | 0.77 ± 0.02 | 0.78 ± 0.01 | 659 |
| Average | 0.78 ± 0.04 | 0.67 ± 0.09 | 0.77 ± 0.04 | **<u>0.81 ± 0.06</u>** | |

**Note:** Underlined bold font face indicates the best performance for each species. \* and \*\* indicate statistical significance compared to all other methods using the Wilcoxon signed-rank test, $p < 0.05$ or 0.01 respectively.

This experimental setting reflects realistic genomic contexts, where complete proteomes are to be annotated. Given the broad diversity of microorganisms and the complexity of their protein sequences, this evaluation provides insight into the models’ ability to handle genome-wide predictions effectively.

### 3.4 Model interpretation for trustworthy predictions

We validated the domain-specific evidence derived from the relevance scores computed by the final HIT-EC model. For the assessment, we compared the relevance scores with characterized motif sites within enzyme sequences of the Cytochrome P450 (CYP) family, specifically CYP106A2 [EC 1.14.15.8]. CYP106A2 functions as a bacterial steroid hydroxylase, capable of hydroxylating various steroids, and its protein structure has also been extensively studied [34]. We considered thirteen protein sequences of the well-characterized bacterial CYP106A2 enzyme family: two from the PDB and eleven from Swiss-Prot, all sharing over 90% sequence similarity with 5XNT [35] and 4YT3 [36]. Then, we computed the relevance scores for each protein sequence, illustrated using multiple sequence alignment (MSA) (e.g., Clustal Omega [37]). We compared the relevance scores of HIT-EC with ECPICK’s ones, as ECPICK was the only approach that provided evidential scores for amino acids.

The interpretation results showed that HIT-EC enhanced the detection of key motif sites (oxygen-binding, EXXR, and heme-binding motifs). Fig. 4 illustrates the CYP106A2 sequences, with functional domains emphasized using colored boxes. Conserved sequences are highlighted in red using ESPript3 [38]. Motif sites, including oxygen-binding, EXXR, and heme-binding domains, are outlined with blue boxes, while substrate recognition sites (SRS 1-6) within the CYP106A2 family are marked with green boxes. The signature regions of the enzyme within the CYP106A2 family are illustrated by purple boxes. The relevance scores of ECPICK and HIT-EC are visualized in Fig. 4 using a color scale, such that high scores are in red and low scores are in white. Both models effectively located the key motif sites within the CYP106A2 family with high scores, including the oxygen-binding motif and heme-binding domain, which are pivotal for defining the first and second levels of EC classification (e.g., 1.14). However, HIT-EC produced more discriminative scores for critical motif regions, including the oxygen-binding site, EXXR motif, and heme-binding motif, compared to ECPICK. Significantly, HIT-EC clearly identified the EXXR motif, which ECPICK did not recognize.

**Fig. 4.**
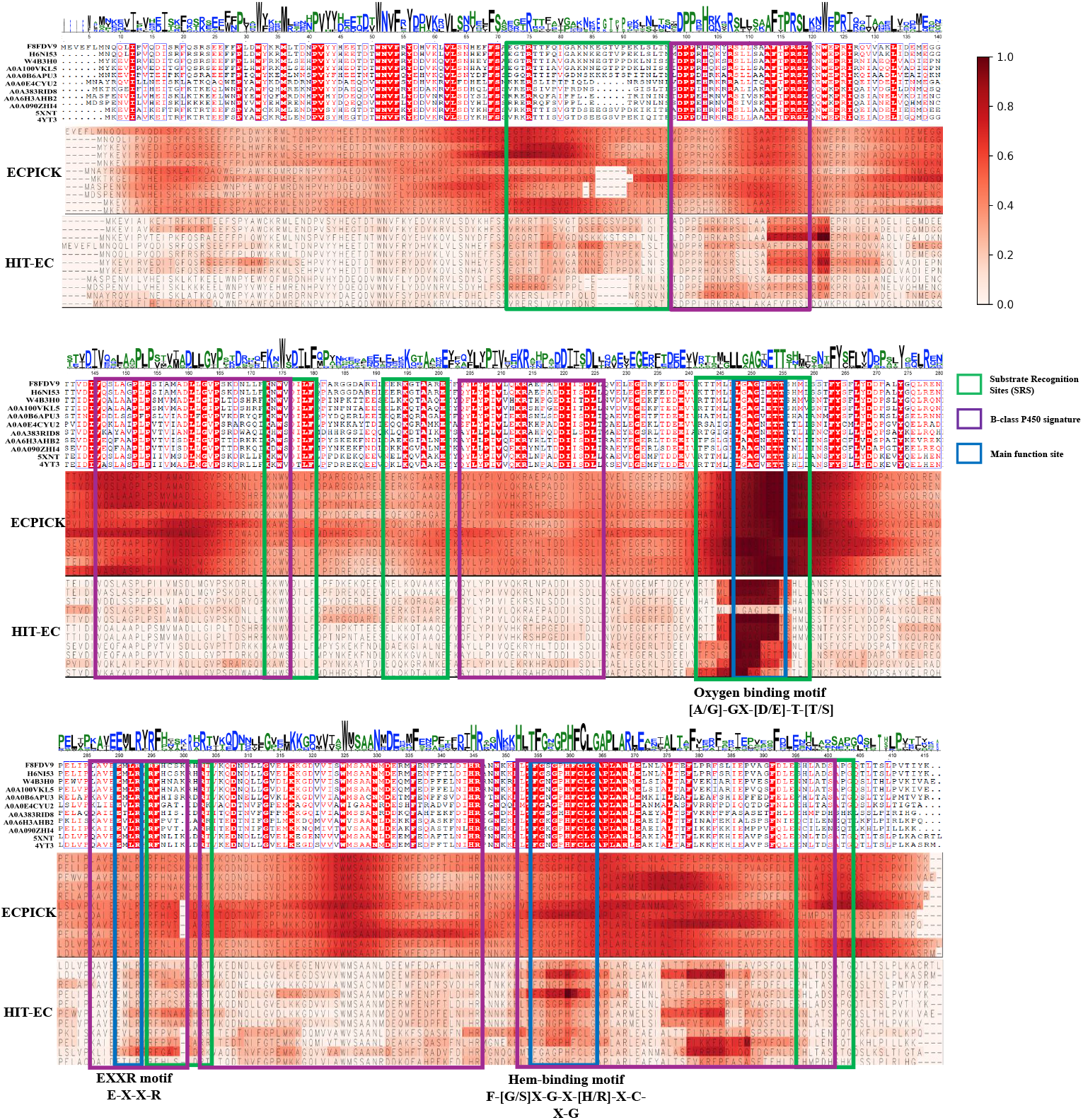
Model interpretation for trustworthy prediction. HIT-EC identifies key amino acids contributing to the prediction, specifying the critical regions essential for the enzyme’s activity as domain-specific evidence for the trustworthy predictions. The blue box corresponds to the main function site of enzymes, the purple box represents the signature region of the enzyme, and the green box denotes the Substrate Recognition Sites (SRS).

## 4 Discussion

In this study, we demonstrated that HIT-EC significantly advances EC number prediction through its innovative hierarchical interpretable transformer approach. HIT-EC achieved superior predictive performance in various evaluation settings, including cross-validation with large datasets, external validation, and species-specific evaluation. Importantly, HIT-EC significantly outperformed the benchmark models in predicting underrepresented EC numbers, which can be attributed to its innovative architectural design and novel learning strategy that handles incomplete annotations. HIT-EC’s transformer-based hierarchical framework captures complex dependencies between the four levels of the EC hierarchy, while preserving broader contextual information of the protein sequence, which results in enhanced predictive power for overall and underrepresented EC classes.

Furthermore, we enhanced the trustworthiness in the predictions through evidential deep learning. By incorporating attention mechanisms and relevance propagation, HIT-EC produces domain-specific evidence for the prediction, which aligns with established biological knowledge. HIT-EC accurately identifies conserved motifs and functional regions in enzymes, as demonstrated in the CYP106A2 family analysis. The evidential approach not only ensures the reliability of its predictions, but also highlights regions of potential biological significance, offering insights that could guide further research and experimental validation.

As a future research direction, further improvement of the predictive power for underrepresented EC numbers would be critical. HIT-EC, while superior to state-of-the-art models, still showed low-performance when classifying underrepresented EC numbers (micro-averaged F1-score of 0.77). Additionally, the computational cost of the hierarchical transformer architecture may pose challenges for large-scale deployments. Future work could focus on optimizing the model’s efficiency without compromising its predictive and interpretative capabilities. HIT-EC represents a significant step forward in EC number prediction, advancing deep learning techniques with interpretability to deliver accurate and trustworthy results.

## Acknowledgments

We acknowledge the supports from National Science Foundation Major Research Instrumentation (NSF MRI) (Grant#:2117941), Ministry of Oceans and Fisheries in the Republic of Korea (20200610, KOPRI Grant), and National Research Foundation of Korea (NRF) by the Korea government (MSIT) (RS-2024-00354012).

## Code availability

The open-source code is publicly available at: https://github.com/datax-lab/HIT-EC.

